# On the inference speed and video-compression robustness of DeepLabCut

**DOI:** 10.1101/457242

**Authors:** Alexander Mathis, Richard Warren

## Abstract

Pose estimation is crucial for many applications in neuroscience, biomechanics, genetics and beyond. We recently presented a highly efficient method for markerless pose estimation based on transfer learning with deep neural networks called DeepLabCut. Current experiments produce vast amounts of video data, which pose challenges for both storage and analysis. Here we improve the inference speed of DeepLabCut by up to tenfold and benchmark these updates on various CPUs and GPUs. In particular, depending on the frame size, poses can be inferred offline at up to 1200 frames per second (FPS). For instance, 278 × 278 images can be processed at 225 FPS on a GTX 1080 Ti graphics card. Furthermore, we show that DeepLabCut is highly robust to standard video compression (ffmpeg). Compression rates of greater than 1,000 only decrease accuracy by about half a pixel (for 640 × 480 frame size). DeepLabCut’s speed and robustness to compression can save both time and hardware expenses.

## I. INTRODUCTION

Pose estimation is an integral part of behavioral analysis [1–3]. DeepLabCut is built around the feature detectors of DeeperCut, one of the best performing pose-estimation algorithms for human poses on challenging benchmarks [4, 5]. Recently, we demonstrated that a small number of training images can be sufficient to train DeepLabCut to within human-level labeling accuracy for various pose-estimation problems in the laboratory [6].

Modern behavioral neuroscience experiments can produce large quantities of data [2, 3, 7–9]. Analyzing and storing video files can consume both time and disk space. Here we address these issues by improving DeepLabCut’s inference speed and demonstrating its robustness to high levels of video compression. This allows users to analyze data faster and store videos much more efficiently (even before) extracting poses. We demonstrate these improvements using two distinct data sets across various video dimensions and on different hardware (Central and Graphical Processing Units [CPUs] and [GPUs]). We hope that this will provide helpful insights and methods for the analysis of large-scale behavioral experiments.

The faster inference code will be integrated with the GitHub-DeepLabCut repository [10]. Example applications ranging from horse gait analysis to mouse whisker tracking can be found at the DeepLabCut website [11]. To use the faster inference, a user only needs to change the *batchsize* parameter when analyzing videos.

## II. RESULTS

We considered videos of two different mouse behaviors: locomotion on a transparent treadmill [12] and odor-guided navigation on a paper spool (trail-tracking) [6]. These videos differed in their dimensions, number of frames, and number of tracked features (554×554 pixels, 10,000 frames, and 14 features for treadmill locomotion; 640×480 pixels, 2,330 frames, and 4 features for trail-tracking). In order to assess the effects of frame size on speed, we downsampled the videos to four different spatial scales (Fig. 1A). We trained one network for each behavior at all scales simultaneously (see methods) and uses those networks benchmark various conditions. We only benchmarked DeepLabCut based on a ResNet-50 [13]. The larger ResNet-101 can be more accurate, but will also be slower [6].

**FIG. 1.**
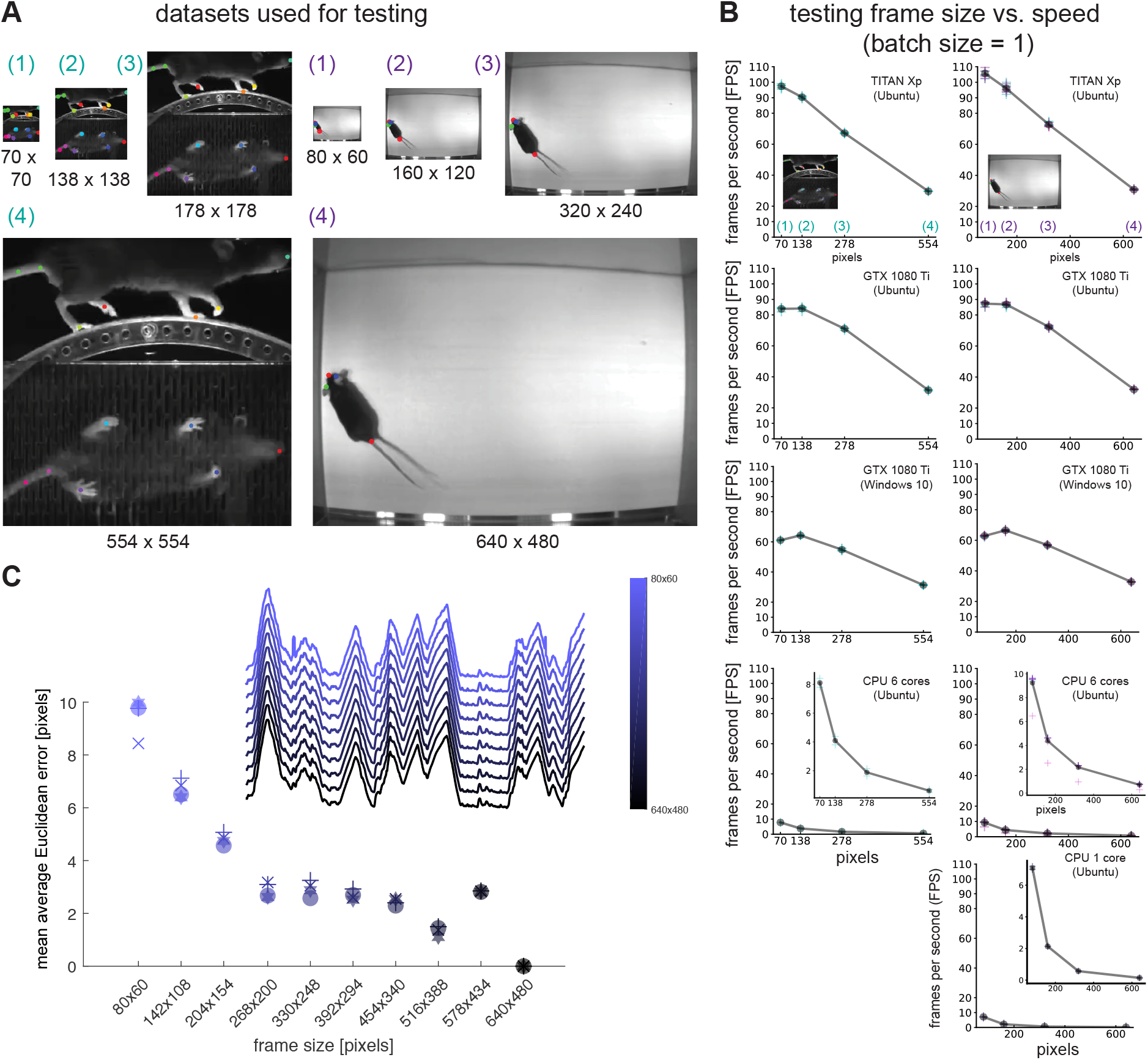
Analysis speed of DeepLabCut for varying video dimensions. **A:** Illustration of the two data sets. We downsampled the largest data to various sizes. For each data set one network was trained at all scales simultaneously. **B:** Inference speed of (original) DeepLabCut for the two data sets as a function of frame width evaluated on various hardware components. Two different trail-tracking videos were considered (cyan and magenta for trail-tracking), as well as one for locomotion (cyan). Each inference was performed 5 times. Average processing speed is depicted in gray. Smaller frames can be processed substantially faster, and GPUs are much faster than CPUs. Multi-core processing only mildly accelerates processing times. All x axes refer to video width. **C:** Inset: Inferred snout x-coordinate vs time for varying video dimensions for trail-tracking data for one example video (data rescaled from 640×480 and predictions are scaled back). The inferred snout position is highly robust to downsampling. We quantified this with the mean average Euclidean error, which is computed by comparing the inferred poses for each body parts against the reference trajectory extracted from 640 × 480 video. The error (per body part) is plotted against video dimensions; snout, left ear, right ear, tail base errors are depicted as disk, +, x and hexagon,respectively. The inferred body positions are robust to downsampling.

### A. Inference speed for DeepLabCut

First, we benchmarked inference speed on videos with varying frame size on 5 different hardware components. As expected, processing time increased with larger frame sizes (Fig. 1B), and inference speed was about 15-60 times faster on GPUs than single CPUs. The multi-core configuration on the CPU only mildly improved performance.

Surprisingly, downsampling images had minimal impact on pose estimation accuracy, with large (¿4 pixels) errors only emerging when scales were reduced to about 30% of their original size (Fig. 1C). Given that smaller images are also processed more quickly (Fig. 1B), this means that analysis speed can be increased by using lower resolution videos without greatly compromising accuracy.

### B. Inference speed for updated DeepLagbCut

We improved the speed of DeepLabCut by 1) allowing the simultaneous processing of multiple frames, and 2) parallelizing the final step in the pose estimation process. Predicted poses are calculated from the scoremap and location-refinement layers by extracting the maximum scoremap locations and shifting them according to the corresponding location refinement predictions [4, 6]. This can quite naturally done by looping over the body parts and frames per batch (double-loop method). We optimized this step by parallelizing it with Numpy [14]. This substantially improved inference speed without affecting result accuracy, particularly when processing multiple frames simultaneously in batches (Fig 2). Thus, we only report inference speeds for the Numpy-based method.

**FIG. 2.**
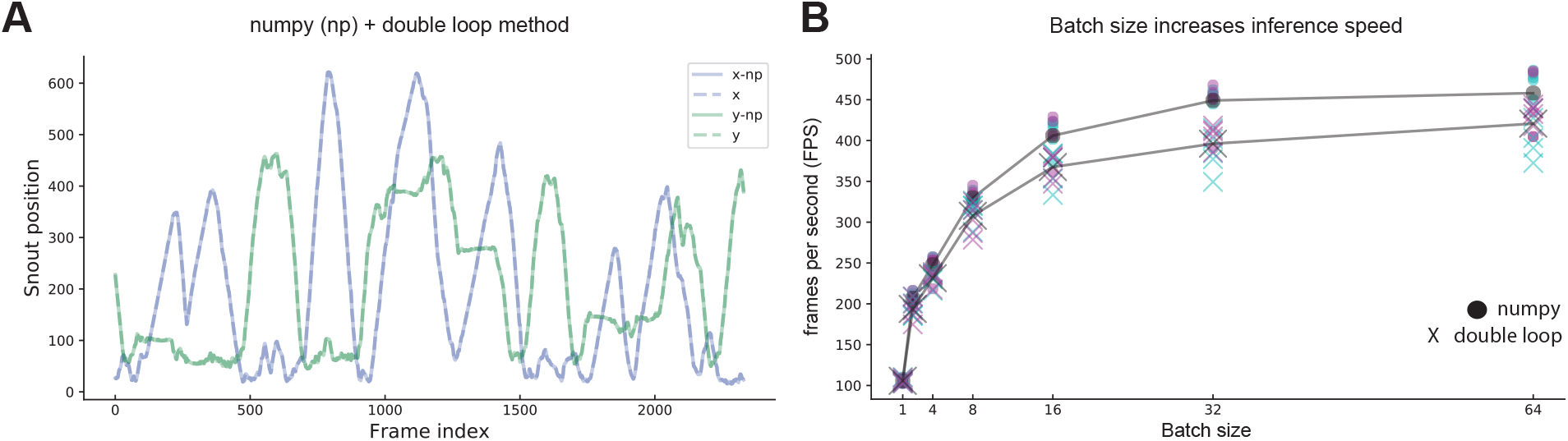
Analysis speed of updated DeepLabCut for varying batch sizes. **A:** Inferred trajectory of snout for one of the trail-tracking test videos with 2,330 frames. The numpy and double-loop inference methods match. **B:** Inference speed vs. batch size for the two methods on the trail-tracking data. As previously cyan and magenta depict individual repetitions of different videos and gray the average. For this test video of 80 × 60, both methods are substantially faster with larger batch sizes reaching processing times of up to 450 FPS. The Numpy method consistently outperforms the double loop implementation.

Next, we benchmarked this updated inference code for the data described in Figure 1. Analyzing frames in batches improved inference speed for the trail-tracking dataset up to five-fold depending on the hardware and video dimensions (Fig. 4). Speed improvements of up to ten-fold were observed for the locomotion data set, which has more tracked body parts, square frame sizes, and different image statistics (Fig. 5). We observed that processing speed is not constant for a given video, but tends to ramp up in the beginning of evaluation (data not shown). Thus, the larger increase in speed for the locomotion dataset might be due to the longer video length (10,000 vs. 2,330 frames).

**FIG. 3.**
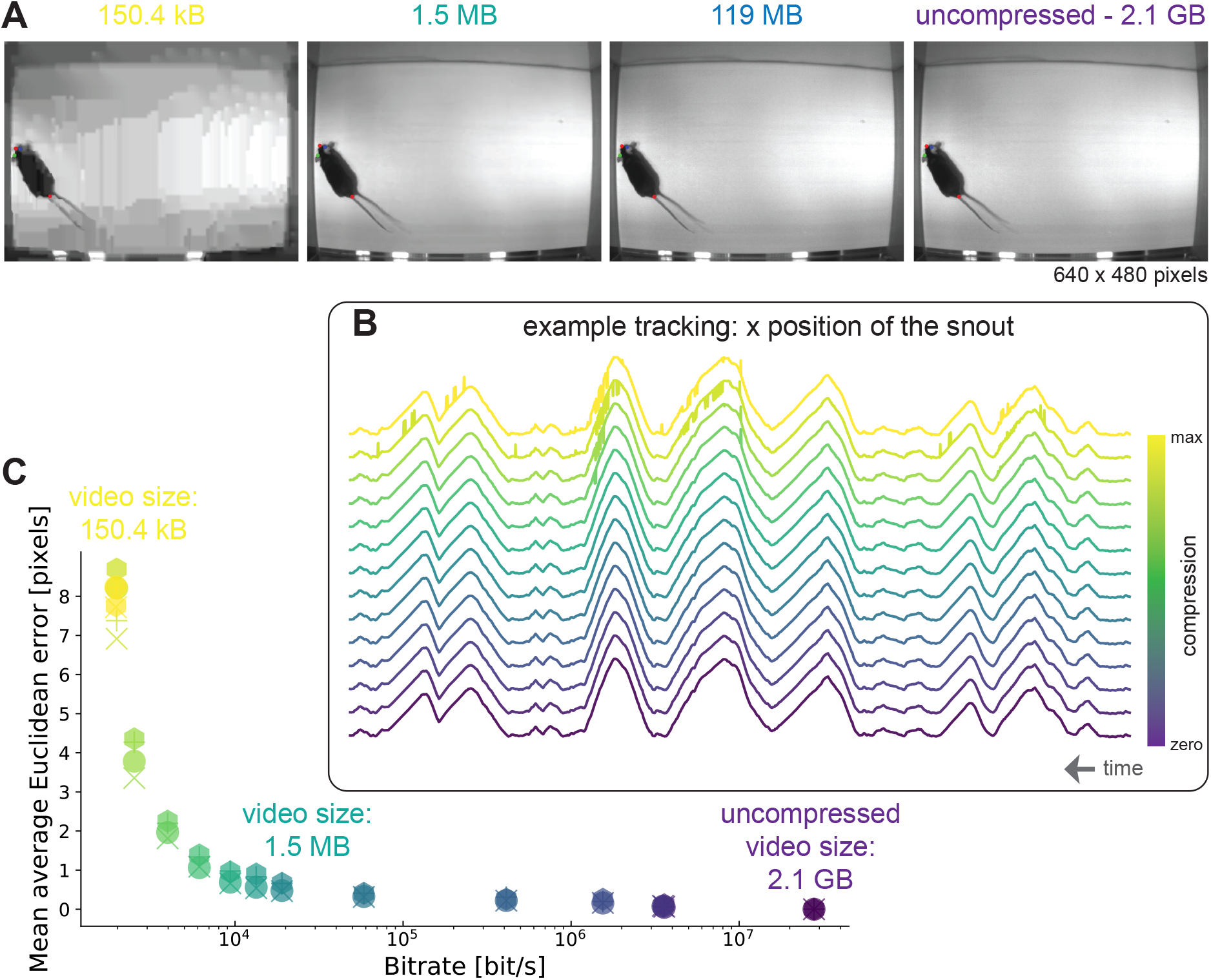
Robustness to ffmpeg (H.264 codec) compression of DeepLabCut: **A:** First frames from example compressed and uncompressed videos with corresponding size listed. The predicted body parts are also depicted. **B:** Overall we considered 13 compression rates. Inferred snout x-coordinate vs time for varying level of video compression. The inferred snout position is highly robust and only starts to jitter for the two most compressed videos. **C:** The mean average Euclidean error is computed by comparing the inferred poses for each body parts against the reference trajectory extracted from the uncompressed video. The error (per body part) is plotted against the bitrate of the video; snout, left ear, right ear, tail base errors are depicted as disk, +, x and hexagon,respectively. All inferred body positions are highly robust when compressed by 3 orders of magnitude. Even beyond that the error only goes up to 8 pixels, which is less than 2% of the frame size.

**FIG. 4.**
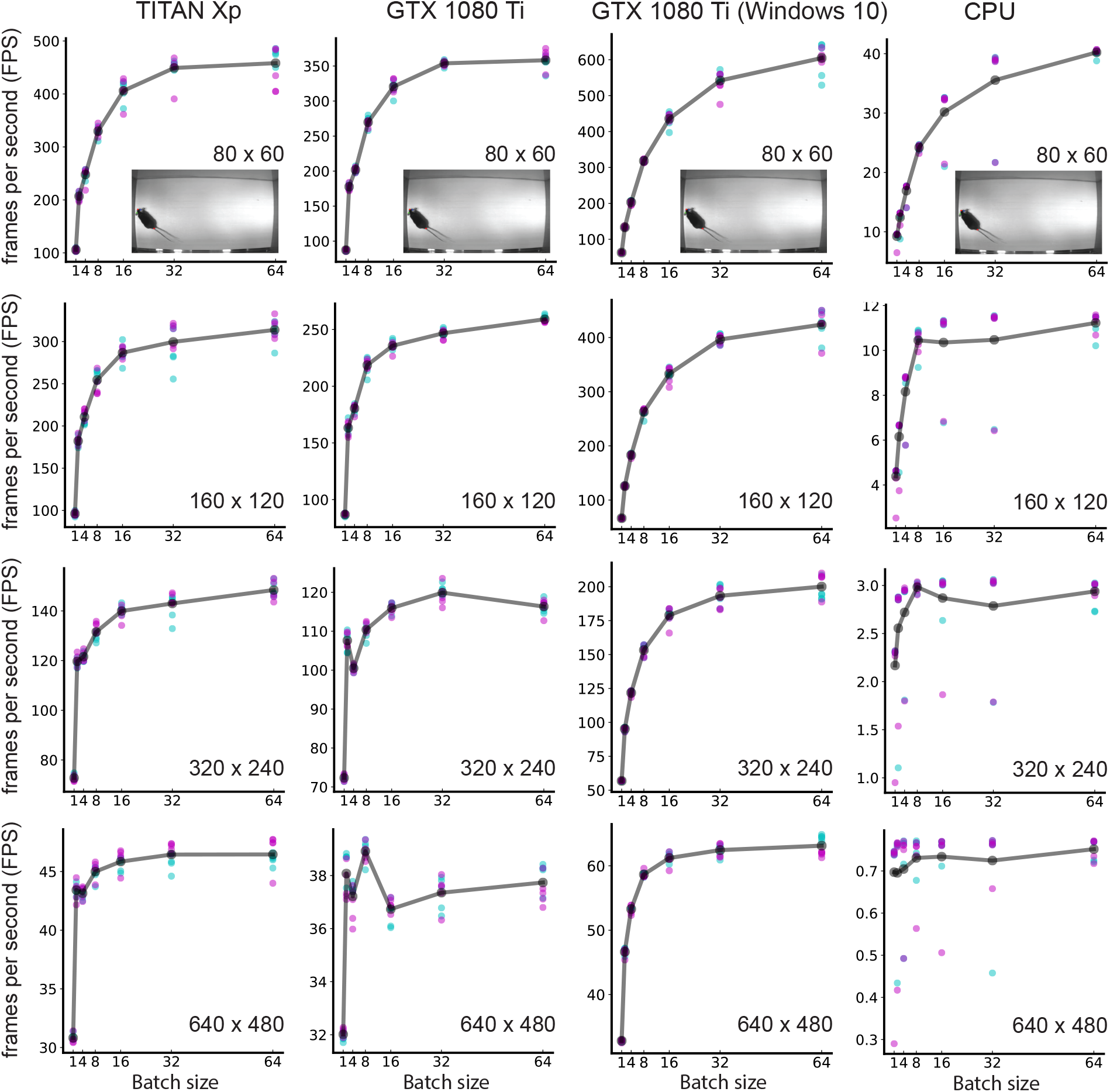
Inference speed of DeepLabCut depending on video dimensions for various computational architectures on the trail-tracking data. Two videos with 2,330 frames were used (magenta and cyan) and ran 5 times in pseudorandom order for each set of conditions. The average inference speed is depicted in gray. Batch processing improved performance by 50% to 500% mostly depending on the frame dimensions. GPUs are 10 − 100 times faster than six-core CPU. Individual GPUs vary widely depending on card and system.

**FIG. 5.**
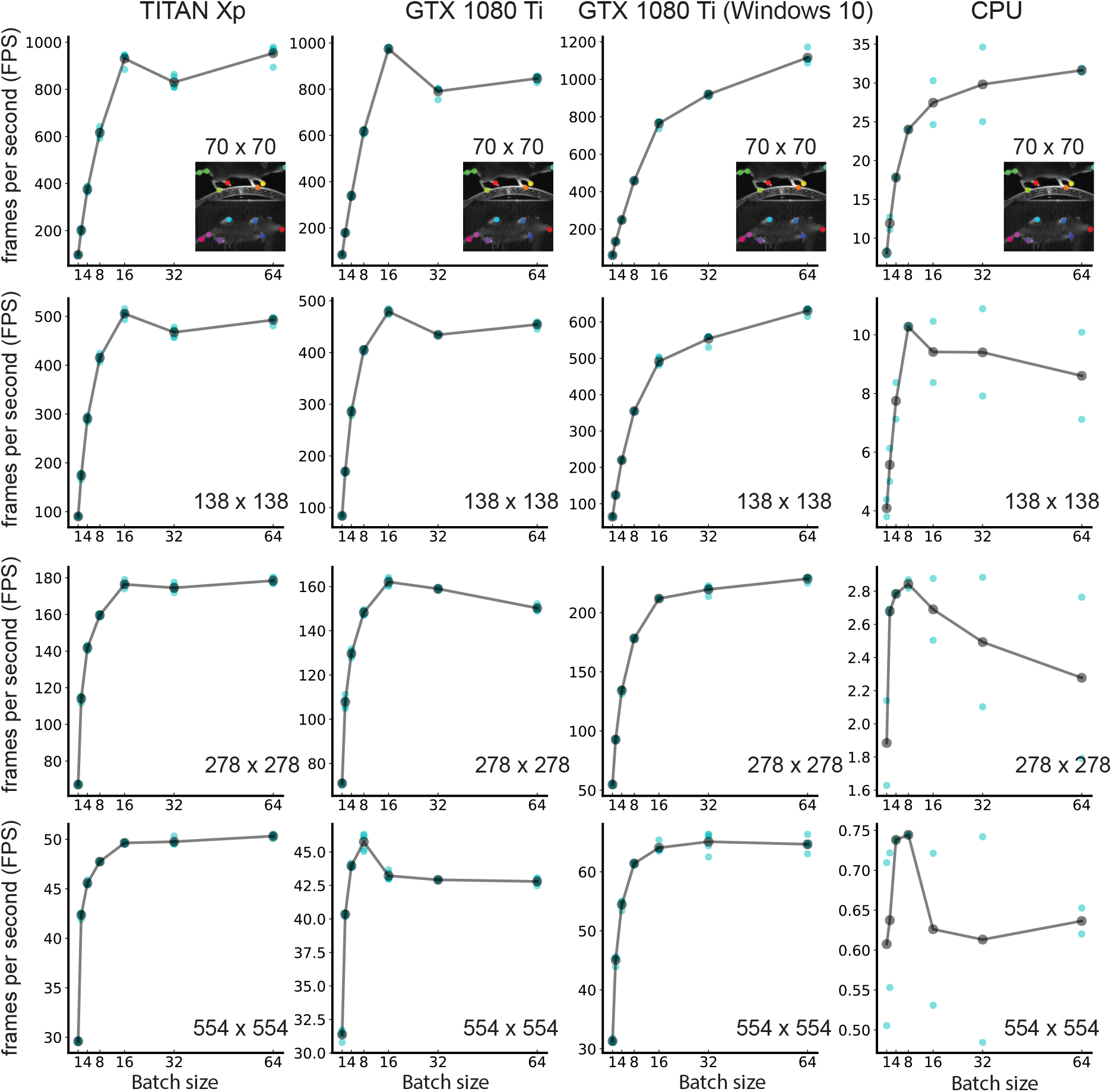
Inference speed of DeepLabCut depending on video dimensions for various computational architectures on locomotion data. One video with 10,000 frames was used (cyan) and ran 5 times in pseudorandom order for each set of conditions. The average inference speed is depicted in gray. Batch processing improved performance by 50% to 500%. GPUs are 10 − 100 times faster than six-core CPU (for time reasons only two iterations were carried out for CPU). Individual GPUs vary widely depending on card and system.

### C. Robustness to standard video compression methods

Modern neuroscience experiments can produce large quantities of data [3, 7–9]. In particular, videos of animal behavior often need to be captured with high frame rates and dimensions to achieve requisite temporal and spatial precision. While video compression [15, 16] can greatly reduce memory demands, it is necessary to ensure videos can be analyzed with comparable accuracy to the original, uncompressed data.

We find that DeepLabCut is remarkably robust to video compression. We compressed the trail-tracking videos by gradually changing the constant rate factor for the FFmpeg H.264 codec (see methods) [17]. We thereby varied the file size from 2.1GB (uncompressed video) to 150.4 kB, which greatly affected video quality (Fig. 3A). We analyzed these videos using the same network as above, trained across all scales and using only frames from uncompressed videos. Remarkably, accuracy remained high even when compressing video sizes 10,000 fold (Fig. 3A). However, severe compression caused extracted postures to jitter over time (Fig. 3B), leading to an average distance for all body parts of around 8 pixels. While this is still quite precise considering the video quality, compression by more than 1,000 only mildly affects accuracy (with less than 1 pixel average error). For comparison, the human accuracy to label body parts was around 3 pixels [6].

These experiments may underestimate the accuracy achievable using highly compressed videos. We trained a network on uncompressed videos but evaluated on compressed videos. Increased accuracy can likely be achieved by training and evaluating on videos compressed in the same manner. Second, the jitter observed at lower bitrates could likely be remedied with simple temporal filtering, e.g. a median filter, as here poses are extracted on a frame-by-frame basis. We did observe that the training across multiple scales improves the robustness to video compression.

## III. CONCLUSIONS

Neuroscience has arrived in the era of big data [7–9, 18]. Being able to analyze and store data efficiently will be critical as datasets become larger and analyses become more complex. We addressed these issues in the context of high-throughput behavioral analysis by improving the speed of DeepLabCut and determining that it is robust to high levels of video compression. Recently another method, called LEAP, using Deep Neural Networks for pose estimation in animals was proposed. LEAP uses a shallow, 15 layer network that makes it fast (185 FPS on 192 × 192 pixel images, but it is not reported how the speed of LEAP scales with frames size and feature number). [19]. Because the weights are trained from scratch, this architecture comes at the price of having to align the animals prior to training and evaluation, which also takes time that is not accounted for. DeepLabCut does not require aligned data and learns efficiently due to transfer learning.

Now DeepLabCut can be used to analyze millions of frames in a matter of hours, all while maintaining modest file sizes. For example, one of us typically collects about 1 million 400×400 pixel frames per day, which would amount to 160 GB for 8 bit uncompressed video. However, compressing online during acquisition results in files that only occupy about 5 GB of disk space that can then be analyzed on a GTX 1080 Ti in approximately 3 hours.

This work offers several practical insights for users. File sizes can be reduced by either compressing videos or decreasing their spatial resolution while minimally impacting analysis accuracy. Furthermore, deceasing spatial resolution can improve analysis speed, again with minimal impact on accuracy. We recommend that users experiment with different compression rates and spatial resolutions to find the right balance between analysis speed, file sizes, and accuracy.

Modern compression standards like H.264 are broadly available with highly optimized implementations across platforms [15, 16]. This makes them highly practical for large scale experiments in combination with DeepLabCut, as we showed here. However, we expect that there will be much progress due to the active development of newer standards and methods [20–23].

## METHODS

**Computer Hardware:** We considered 5 different hardware scenarios: NVIDIA GTX 1080Ti and NVIDA Titan Xp with TensorFlow 1.2 run inside a Docker container; two CPU cases with TensorFlow 1.8 (an Intel Xeon CPU E5-2603 v4 @ 1.70GHz) either with all 6 cores or just 1 engaged. These four cases were tested on an Ubunutu 16.04 computer. Additionally, we utilized a NVIDIA GTX 1080 Ti with TensorFlow 1.8.0 on Windows 10 with Intel i7-8700K.

**Trail-tracking data set:** All surgical and experimental procedures for trail-tracking were in accordance with the National Institutes of Health Guide for the Care and Use of Laboratory Animals and approved by the Harvard Institutional Animal Care and Use Committee. Experiments were carried out in the laboratory of Prof. Venkatesh N. Murthy at Harvard University. We took 511 of the low-resolution camera images from the olfactory trail-tracking data in DeepLabCut [6]. Those frames have 640 × 480 pixels and were recorded with a Point Grey Firefly (FMVU-03MTM-CS) camera. To train networks with different scales we downsampled those frames by factors (of 1), 2, 4 and 8. Overall we trained one network with the same train/test split (of images) for all scales. Furthermore, we utilized two different test videos that contained 2,330 test frames each. We evaluated each video 5 times for each hardware component.

**KineMouse Wheel data set:** All surgical and experimental procedures for trail-tracking were in accordance with the National Institutes of Health Guide for the Care and Use of Laboratory Animals and approved by the Columbia Institutional Animal Care and Use Committee. Experiments were carried out in the laboratory of Prof. Nathaniel B. Sawtell at Columbia University. Details on how to construct the wheel have been posted at Hackaday [12]. Videos were taken with a single Point Grey Grasshopper3 (GS3-U3-23S6C-C) camera and were compressed during acquisition using ffmpeg and Bonsai acquisition software [24]. Top and bottom views were cropped from a single camera online during acquisition and concatenated offline to yield 396 × 406 pixel videos. One 10,000 frame video was resized to 554×554 (to approximately match the total number of pixels in the trail-tracking dataset). Training and downsampling were then conducted as described for the trail-tracking data set.

**Benchmarking Analysis:** Beyond the hardware components (see above), we varied the batch size {1, 2, 4, 8, 16, 32, 64}, video dimensions (depending on data set), analysis method (standard, double-loop, and Numpy), and performed 5 repetitions for each set of conditions. Thereby, “standard” refers to the original DeepLabCut method for batch size 1. For each experimental data set and hardware configuration we evaluated the conditions in pseudorandom order to minimize influences of the state of the hardware. However, given that the experiments ran for several hours (GPU) or days (CPU) this means that the hardware was engaged.

**Compression Analysis:** For the robustness to compression results, we converted one uncompressed video in the command line by varying `factor`:
`ffmpeg-i videoinput.avi-vcodec libx264-crf factor videocompressed.mp4`

We considered the following factor parameters: {100, 50, 45, 40, 35, 30, 27, 25, 20, 15, 10, 1, 0.1}. This led to a videos ranging from 2.1GB (uncompressed) to around 150kB. The compression was done with ffmpeg version 2.8.15-0ubuntu0.16.04.1.

**Code:** The Code for DeepLabCut can be found on our GitHub repository: https://github.com/AlexEMG/DeepLabCut. Examples of analyzed behaviors can be found on the DeepLabCut website.

## Acknowledgments

We are grateful to Mackenzie Mathis for help preparing the figures and comments on the manuscript, as well as members of the Bethge, Dulac, Murthy, Mathis and Sawtell labs for discussions.

